# Inherited retinal degenerations: PARP regulates calpain activation via TRPM2 channels in *rd1* mouse photoreceptors

**DOI:** 10.1101/2025.11.13.688383

**Authors:** Jie Yan, Lei Kong, Zhijian Zhao, QianLu Yang, Lan Wang, QianXi Yang, Christian Harteneck, Kangwei Jiao, Zhulin Hu, François Paquet-Durand

## Abstract

Inherited retinal degeneration (IRD) refers to untreatable blinding diseases characterized by progressive photoreceptor loss. Photoreceptor degeneration is often associated with an excessive activation of poly (ADP-ribose) polymerase (PARP) and Ca^2+^-dependent calpain-type proteases. To explore the interplay between PARP and calpain activity, we employed organotypic retinal explant cultures derived from wild-type mice and from the *rd1* mouse model for IRD. Retinae were treated with the PARP inhibitors INO1001 or Olaparib, the poly (ADP-ribose) glycohydrolase (PARG) inhibitor JA2131, or the transient receptor potential channel M2 (TRPM2) blocker 8-Br-ADPR.

Readouts included the TUNEL assay to detect cell death, *in situ* activity assays for histone-deacetylases (HDAC), PARP, and calpain, as well as immunostaining for activated calpain-2, and poly (ADP-ribose) (PAR). PARP, PARG, and TRPM2 inhibition reduced calpain activity and calpain-2 activation. PARP activity was decreased by PARP and TRPM2 inhibitors but not by PARG inhibition. Remarkably, the PARP inhibitor INO1001 increased HDAC activity unlike any of the other compounds. When combined with the PARG inhibitor JA2131, INO1001 reduced photoreceptor cell death in a synergistic fashion, although such synergy was not observed for calpain or PARP activity. Moreover, synergistic photoreceptor preservation was not observed when JA2131 was combined with the PARP inhibitor Olaparib.

Overall, these results indicate that in *rd1* photoreceptors, PARP controls calpain activity via PARG and TRPM2-induced Ca^2+^ influx. We also characterize INO1001 as potentially more beneficial for IRD treatment than Olaparib. Our study details the complexity of PARP-signalling in photoreceptors and identifies PARG and TRPM2 as new targets for IRD therapy development.

## INTRODUCTION

Inherited retinal degeneration (IRD) represents a diverse group of progressive, neurodegenerative diseases that often lead to blindness. IRD is characterized by progressive photoreceptor cell death [1] and is usually untreatable due to its genetic heterogeneity and lack of effective therapies. Within the IRD group the most common disease is *retinitis pigmentosa* (RP) affecting approximately two million people all over the world [2]. Early stage patients experience a loss of night vision due to the primary degeneration of rod photoreceptors. At late stages the patient’s field of vision narrows until only central vision remains (*i.e.* “tunnel vision”). Eventually, the secondary degeneration of cone photoreceptors results in total blindness [2]. The second messenger cyclic-guanosine-monophosphate (cGMP) has been found to play a central role in the pathobiology of many genetically distinct types of IRD. In photoreceptor cells excessive cGMP-signalling may be directly or indirectly related to the activity of poly (ADP-ribose) polymerase (PARP), Ca^2+^-dependent, calpain-type proteases and histone deacetylase (HDAC) [3–5].

The *rd1* mouse (retinal degeneration 1), is a laboratory mouse strain commonly used in research on retinal degeneration. It is a naturally-occurring genetic IRD model first described by Keeler one century ago [6]. The *rd1* mouse carries a mutation affecting the expression of the beta subunit of phosphodiesterase 6 (PDE6). Since in rod photoreceptors PDE6 beta is responsible for the hydrolysis of cGMP, the *rd1* mutation leads to an accumulation of cGMP and rapid rod loss [7]. In photoreceptors cGMP is known to act upon two prototypic targets: 1) the cyclic-nucleotide-gated (CNG) channel, which allows for Ca^2+^-influx and is the main effector of the phototransduction cascade, and 2) protein kinase G (PKG), which phosphorylates a large number of target proteins [8]. Probably further downstream from these two cGMP targets, cGMP accumulation in *rd1* mouse rods is associated with a prominent activation of PARP, calpain, and HDAC [9].

PARP is an enzyme family involved in DNA repair processes. When detecting DNA damage, PARP is activated and binds to the damaged sites, where it facilitates the recruitment of other DNA repair proteins to the site of damage [10]. PARP uses nicotinamide adenine dinucleotide (NAD^+^) as a substrate to synthesize and attach poly (ADP-ribose) (PAR) chains onto itself and other target proteins. PAR polymers help to recruit and activate other DNA repair factors, creating a scaffold for efficient repair processes [11]. Covalently attached PAR can be hydrolysed by poly (ADP-ribose) glycohydrolase (PARG) to free PAR polymers or to mono adenosine diphosphate ribose (ADP-ribose, ADPR) [12]. Paradoxically, excessive activation of PARP may also drive a specific form of cell death, termed PARthanatos, a phenomenon that has been linked to excessive NAD^+^ consumption or PAR generation [13].

ADPR resulting from PARP activity is the main activator for transient receptor potential cation channel subfamily M member 2 (TRPM2) [14]. TRPM2 belongs to the transient receptor potential (TRP) superfamily of ion channels [15]. It is a Ca^2+^-permeable channel that is primarily expressed in immune cells, neurons, and various other tissues. Activation of TRPM2 channel leads to an influx of Ca^2+^ ions into the cell, which, in turn, may trigger downstream signalling cascades and cellular responses [16].

For instance, TRPM2 mediated Ca^2+^-influx can lead to the activation of Ca^2+^-dependent, calpain-type proteases that play important roles in various cellular processes such as cell signalling, cytoskeletal remodelling, cell migration, and programmed cell death [17]. When Ca^2+^ binds to the Ca^2+^-binding domain within calpain enzymes, it triggers their activation and enables them to selectively cleave target proteins at specific sites [18].

HDAC regulates gene expression by removing acetyl groups from histones, impacting chromatin structure and gene activity [19]. HDACs are categorized into two main groups based on their cofactor dependency: 1) Zinc-dependent HDACs: These include Class I, II, and IV HDACs. Class I HDACs (HDAC 1, 2, 3, 8) are primarily found in the nucleus and resemble yeast RPD3 deacetylase. Class II HDACs are further divided into IIa (HDAC 4, 5, 7, 9) and IIb (HDAC 6, 10), which shuttle between the nucleus and cytoplasm. Class IV comprises only HDAC 11, which shares features with both Class I and II. 2) NAD^+^-dependent HDACs: known as Class III or Sirtuins, these HDACs require NAD^+^ to function and differ significantly in mechanism and function from the zinc-dependent classes [20]. Over-activation of HDAC may trigger photoreceptor cell death [21]. However, neuroprotection exhibited by Sirtuins has sparked debate about the role of HDACs in RP [22, 23].

Our previous research suggested that PARP and calpain take part in the same cell death pathway triggered by excessive cGMP-signalling [24]. PARP appears to control calpain activity while calpain does not regulate PARP activation [9]. Nevertheless, how exactly PARP regulates calpain activity is still unknown. Here, we hypothesized that PARP may control calpain activation via ADPR-induced TRPM2 activation. To investigate this possibility, we used specific inhibitors for PARP, and PARG, as well as an inhibitory ADPR analogue in organotypic *rd1* retinal explant cultures. Through these interventions, we show that (1) PARP, PARG, and ADPR can promote photoreceptor cell death, and that (2) PARP regulates calpain activity through ADPR-dependent activation of TRPM2, and that (3) the PARP inhibitor INO1001 displays superior therapeutic efficacy compared to Olaparib.

## MATERIALS AND METHODS

### Animals

For retinal explant cultures C3H/HeA *Pde6b ^rd1^*^/*rd1*^ animals (*rd1*), their congenic wild-type C3H/HeA *Pde6b*^+/+^ counterparts (*wt*) [25], and B6.129SvJ;C3H/HeA-*CNGB1*^tm^ double-mutant mice (*rd1*Cngb1^−/−^*) were used [26]. The *rd1*Cngb1^−/−^* double mutants were generated by an intercross of *rd1* and *Cngb1*^−/−^ and have been maintained by repeated backcrossing over 10 generations to make a congenic inbred strain, homozygous for both gene mutations. Animals were housed under standard white cyclic lighting, had free access to food and water, and were used irrespective of gender. Animal protocols compliant with §4 of the German law of animal protection were reviewed and approved by the Tübingen University committee on animal protection (Einrichtung für Tierschutz, Tierärztlicher Dienst und Labortierkunde, Registration numbers: AK02/19M, AK01/20M).

### Organotypic retinal explant culture

To study the effects of various drugs on photoreceptor enzyme activities and cell death, *wt*, *rd1*, and *rd1*Cngb1^−/−^* retinas were explanted at postnatal day 5 (P5). The retinal explants were maintained in culture until either P11 (wt, rd1) or P17 (wt, *rd1*Cngb1^−/−^*). The explants were cultured on a polycarbonate membrane (83.3930.040; 0.4 µm TC-inserts, SARSTEDT, Hildesheim, Germany) with complete R16 medium (Gibco; with supplements) [27]. During the cultivation, the complete R16 medium was changed every two days with treatment. The two retinas obtained from a single animal were split across different experimental groups so as to maximize the number of independent observations acquired per animal. Cultures were treated with 0.1 µM INO1001 (IC_50_: ≈ 3 nM [28, 29], A20680; AdooQ, Irvine, USA), 5 µM JA2131 (IC_50_: 0.4 μM [30], HY-137924; MedChemExpress, Sollentuna, Sweden), 50 µM 8-Br-ADPR (IC_50_: 5 μM [31], No.: B 051; BIOLOG, Bremen, Germany), and 1 µM Olaparib (IC_50_: 5 nM [32], T3015; TargetMol; Boston; USA) for monotherapies and 0.1 µM INO1001 and 5 µM JA2131 or 1 µM Olaparib and 5 µM JA2131 for combined therapies (The details of compounds were presented in Table S1). In these treatments, all compounds were dissolved in DMSO at a final medium concentration of no more than 0.1% DMSO. Cultures were ended at P11 (*rd1*), and P17 (*rd1*Cngb1^-/-^*) by either fixation with 4% paraformaldehyde (PFA) or without fixation and direct freezing in liquid N_2_. Explants were embedded in Tissue-Tek (Sakura Finetek Europe B.V., Alphen aan den Rijn, The Netherlands) and sectioned (14 µm) in a cryostat (Thermo Fisher Scientific, CryoStar NX50 OVP, Runcorn, UK).

### TUNEL assay

TUNEL (terminal deoxynucleotidyl transferase dUTP nick end labelling) assay kit (Roche Diagnostics, Mannheim, Germany) was used to label dying cells. Histological sections from retinal explants were dried and stored at −20 °C. The sections were rehydrated with phosphate-buffered saline (PBS; 0.1 M) and incubated with proteinase K (1.5 µg/µL) diluted in 50 mM TRIS-buffered saline (TBS; 1 µL enzyme in 1 mL TBS) for 15 min. This was followed by 3 times 5 min TBS washing and incubation with blocking solution (10% normal goat serum, 1% bovine serum albumin, and 1% fish gelatine in phosphate-buffered saline with 0.03% Tween-20). TUNEL staining solution was prepared using 21 parts of blocking solution, 18 parts of TUNEL labelling solution, and 1 part of TUNEL enzyme. After blocking, the sections were incubated with TUNEL staining solution overnight at 4 °C. Finally, sections were washed 2 times with PBS, mounted using mounting medium with DAPI (ab104139; Abcam), and imaged by microscopy.

### Calpain activity assay

To visualize overall calpain activity *in situ* on unfixed tissue sections, retinal tissue sections were incubated and rehydrated for 15 min in a calpain reaction buffer (CRB) (5.96 g HEPES, 4.85 g KCl, 0.47 g MgCl_2_, and 0.22 g CaCl_2_ in 100 mL ddH_2_O; pH 7.2) with 2 mM dithiothreitol (DTT). Tissue sections were incubated for 3 h at 37 °C in CRB with tBOC-Leu-Met-CMAC (25 µM; A6520; Thermo Fisher Scientific, OR, USA). Then, sections were washed with PBS and incubated with ToPro (1:1000 in PBS, Thermo Fisher Scientific) for 15 min. Afterwards, tissue sections were washed twice in PBS (5 min) and mounted using Vectashield without DAPI (Vector Laboratories Inc., Burlingame, CA, USA) for immediate visualization by microscopy.

### PARP activity and PAR staining

The PARP *in situ* activity assay is based on the incorporation of a fluorescent NAD^+^ analogue and allows resolving the overall PARP enzyme activity on unfixed tissue sections [33]. Such sections were incubated and rehydrated for 10 min in PBS. The reaction mixture (10 mM MgCl_2_, 1mM dithiothreitol, and 50 μM 6-Fluo-10-NAD^+^ (Cat. Nr.: N 023; Biolog) in 100 mM Tris buffer with 0.2% Triton X100, pH 8.0) was applied to the sections for 3 h at 37 °C. After three 5 min washes in PBS, sections were mounted in Vectashield with DAPI (Vector Laboratories) for subsequent microscopy.

For the detection of PAR, we used an immunostaining enhanced by 3,3′-diaminobenzidine (DAB) staining. The procedure is initiated by quenching of endogenous peroxidase activity using 40% MeOH and 10% H_2_O_2_ in PBS with 0.3% Triton X-100 (PBST) in retinal tissue sections for 20 min. Sections were further incubated with 10% normal goat serum (NGS) in PBST for 30 min, followed by anti-PAR antibody (1:200; ALX-804-220-R100; Enzo Life Sciences, Farmingdale, NY, USA) incubation overnight at 4 °C. Incubation with the biotinylated secondary antibody (1:150, Vector in 5% NGS in PBST) for 1 h was followed by the Vector ABC Kit (Vector Laboratories, solution A and solution B in PBS, 1:150 each) for 1 h. DAB staining solution (0.05 mg/mL NH_4_Cl, 200 mg/mL glucose, 0.8 mg/mL nickel ammonium sulphate, 1 mg/mL DAB, and 0.1 vol. % glucose oxidase in phosphate buffer) was applied evenly, incubated for precisely 3 min, and immediately rinsed with phosphate buffer to stop the reaction. Sections were mounted in Aquatex (Merck, Darmstadt, Germany).

### HDAC activity assay

This assay allows detecting overall HDAC activity *in situ* on fixed tissue sections and is based on an adaptation of the FLUOR DE LYS®-Green System (Biomol, Hamburg, Germany). Retinal sections were exposed to 50 µM FLUOR DE LYS®-SIRT1 deacetylase substrate (BML-Kl177-0005; ENZO, New York, USA) with 2 mM NAD^+^ (BML-KI282-0500; ENZO) in assay buffer (50 mM Tris/HCl, 137 mM NaCl; 2.7 mM KCl; 1mM MgCl2; pH 8.0) for 3 h at 37 °C. Sections were then washed in PBS and fixed in methanol at -20 °C for 20 min. Slides were mounted with FLUOR DE LYS® developer II concentrate (BML-KI176-1250; Enzo) diluted 1:5 in assay buffer overnight for subsequent microscopy.

### Immunohistochemistry for calpain-2 and TRPM2

Sections were rehydrated with PBS for 15 min and then incubated with a blocking solution (10% NGS, 1% BSA, and 0.3% PBST) for 1 h. The primary antibodies, rabbit-anti-calpain-2 (1:200; ab39165; Abcam), and rabbit-anti-TRPM2 (1:200; NB110-81601; Novus Biologicals, Colorado, USA) were diluted in blocking solution and incubated overnight at 4 °C rinsing with PBS for 3 times 10 min each; this was followed by incubation with the secondary antibodies, goat-anti-rabbit AlexaFluor488 (1:400; A11034; Molecular Probes; Oregon, USA), and goat-anti-rabbit AlexaFluor568 (1:300; A11036; Molecular Probes) for 1 h. The sections were rinsed with PBS for 3 times 10 min each and mounted with mounting medium with DAPI (Abcam).

### Microscopy and image analysis in retinal cultures

Images of organotypic explant cultures were captured using a Zeiss Imager Z.2 fluorescence microscope, equipped with ApoTome 2, an Axiocam 506 mono camera, and HXP-120V fluorescent lamp (Carl Zeiss Microscopy, Oberkochen, Germany). Excitation (λExc.) and emission (λEm.) characteristics of filter sets used for different fluorophores were as follows (in nm): DAPI (λExc. = 369 nm, λEm. = 465 nm), AF488 (λExc. = 490 nm, λEm. = 525 nm), AF568 (λExc. = 578 nm, λEm. = 602 nm), and ToPro (λExc. = 642 nm, λEm. = 661 nm). The Zeiss Zen 2.3 blue edition software was used to capture images (tiled and z-stack, 20x magnification). Sections of 14 µm thickness were analysed using 12–15 Apotome Z-planes. For quantification of positive cells in the ONL, we proceeded as follows: The number of cells in six different rectangular ONL areas was counted manually based on the number of DAPI-stained nuclei, and used to calculate an average ONL cell size. This average ONL cell size was used to calculate the total number of cells in a given ONL area. The percentage of positive cells was then calculated by dividing the absolute number of cells positive for a given marker by the total number of ONL cells. The signal intensity on retinal sections was quantified by Zeiss Zen software version 3.0 (Carl Zeiss).

### Statistical analysis and software use

Two-way comparisons were analysed using Student’s *t*-test. Multiple comparisons were made using a one-way analysis of variance (ANOVA) test with Tukeýs multiple comparison test. Calculations were performed with GraphPad Prism 8 (GraphPad Software, La Jolla, CA, USA); *p* < 0.05 was considered significant; Levels of significance were as follows: *, *p* < 0.05; **, *p* < 0.01; ***, *p* < 0.001; ****, *p* < 0.0001. Data in figure 5A was normalized by linear scaling according to the formula: χ*_scaled_=*χ*-*χ*_min_/*χ*_max_-*χ*_min,_* using SPSS Statistics 26 (IBM, Armonk, New York, USA). The figures were prepared using Photoshop 2022 and Illustrator 2022 (Adobe, San Jose, CA, USA). Spearman analyses were performed by R software (Version 4.0.1). The diagram in Figure 7 was created with BioRender.com.

## RESULTS

### TRPM2 is differentially expressed in *wt* and *rd1* retina

The *rd1* photoreceptor degeneration begins around post-natal day (P) 9 followed by a significant rise of dying cells in the outer nuclear layer (ONL; *i.e.* the photoreceptor layer) first seen at P11 [34]. By P20 the *rd1* mouse retina had lost nearly all rod photoreceptors and a large fraction of cone photoreceptors. Our previous research had identified a strong up-regulation of PARP together with an accumulation of PAR in dying *rd1* photoreceptors [34]. Furthermore, we had shown that cGMP-dependent photoreceptor degeneration was strongly decreased after a knock-out of the PARG110 isoform [35]. Since ADPR monomers generated by the combined activity of PARP and PARG can activate TRPM2, we hypothesized that TRPM2 could be involved in *rd1* degeneration.

To assess such a possible association of TRPM2 expression with *rd1* degeneration, we first performed an immunostaining using an antibody directed against TRPM2. In the early post-natal retina, at P11,

TRPM2 was found to be expressed in both *rd1* and *wt* retinas, notably in photoreceptor segments, inner nuclear layer, and in retinal ganglion cells (Figure 1A). However, at P11 the relative TRPM2 signal overall appeared to be stronger in *rd1* retinas than in *wt*, an observation confirmed by fluorescence intensity distribution plots (Figure 1B). In adult *wt* retina at P30, TRPM2 co-staining with the cone-specific marker PNA peanut agglutinin (PNA) showed that TRPM2 was expressed in cone outer segments (Figure 1C). Taken together, these results illustrated the expression of TRPM2 protein in the retina, including in photoreceptors.

**Figure 1.**
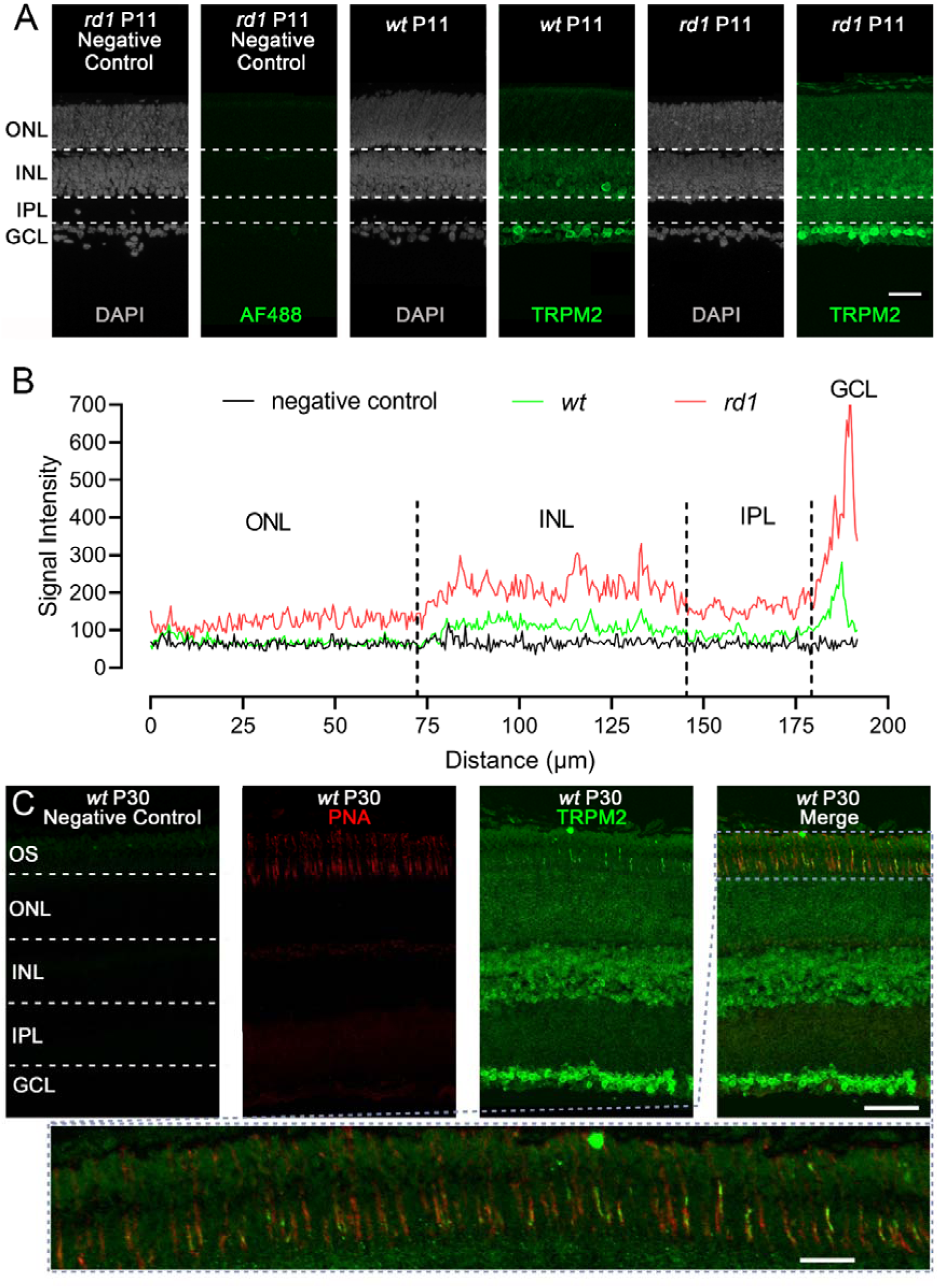
Retinal TRPM2 expression in *wt* and *rd1* retina. **A**) Immunostaining for TRPM2 (green) in post-natal day (P) 11 wild-type (*wt*) and *rd1* retina. DAPI (grey) was used as a nuclear counterstain. **B**) Vertical localisation of TRPM2 signal in *wt* and *rd1* retina illustrated by an intensity distribution plot. Traces show average signal intensities from n=3 *wt* and *rd1* specimens, the red trace indicates the *rd1* situation, green represents *wt*, black represents negative control. **C**) Co-immunostaining for TRPM2 and the cone marker peanut agglutinin (PNA). TRPM2 was found to be partially co-localized with PNA in the outer segments of photoreceptors, indicating it’s expression in cones. Scale bar = 50 µm. OS = outer segment, ONL = outer nuclear layer, INL = inner nuclear layer, IPL = inner plexiform layer, GCL = ganglion cell layer.

### Inhibition of TRPM2 changes the enzymatic activity of PARP

The PARP enzyme uses NAD^+^ to generate PAR polymers which in turn can be degraded by PARG to ADP-ribose (ADPR). ADPR monomers can activate TRPM2 channels, allowing for Ca^2+^-influx (Figure 2A). Calpains are sensitive to intracellular Ca^2+^ levels and among the various calpain isoforms calpain-2 is activated by high, millimolar Ca^2+^ concentrations [18]. Calpain activity and labelling of activated calpain-2 may therefore be used to indirectly determine intracellular Ca^2+^ overload in photoreceptors. To investigate the relationship between TRPM2 and PARP activity as well as between TRPM2 and PARG activity during *rd1* degeneration, we used a range of different compounds to selectively inhibit these cellular targets. Along the putative PARP-signalling pathway (Figure 2A), treatments with the inhibitors INO1001, JA2131, and 8-Br-ADPR were used to inhibit either PARP, PARG, or TRPM2, respectively. The well-established PARP inhibitor Olaparib, which was previously tested in *rd1* retina [36], was used for additional controls. Dose response curves for INO1001, JA2131, and 8-Br-ADPR were generated to select appropriate treatment concentrations (Figure S1A, B, C). For an overview of the compounds used and their inhibitory capacities please refer to Figure S1 and Table S1.

**Figure 2.**
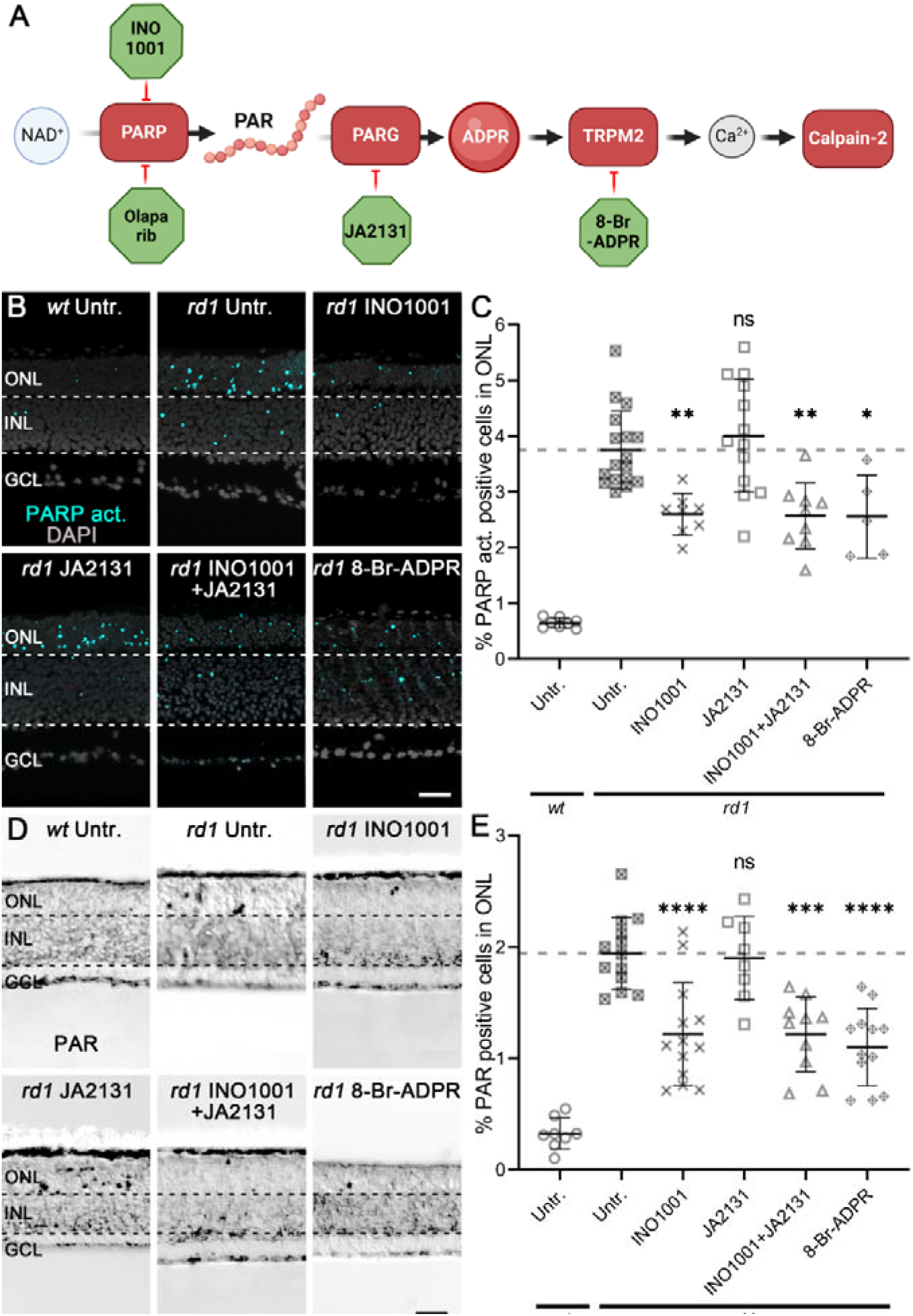
INO1001 and 8-Br-ADPR effectively reduce PARP activity and PAR generation. **A**) Diagram illustrating how PARP-signalling may influence Ca^2+^-influx and calpain-2 activity. PARP, PARG, and TRPM2 were inhibited using INO1001, JA2131, 8-Br-ADPR, respectively. **B**) PARP activity assay (cyan) was performed in *rd1* and wild-type (*wt*) retinal explant cultures. DAPI (grey) was used as nuclear counterstain. Untreated (Untr.) *rd1* and *wt* retina were compared to *rd1* retina treated with either INO1001, JA2131, INO1001 combined with JA2131, or 8-Br-ADPR. **C**) Scatter plot showing percentage of PARP activity positive cells in the outer nuclear layer (ONL). Untr. *wt*: 8; Untr. *rd1*: 17; INO1001 *rd1*: 8; JA2131 *rd1*: 13; INO1001+JA2131 *rd1*: 9; 8-Br-ADPR *rd1*: 5. **D**) PAR staining (black) was performed in *rd1* and *wt* retinal explant cultures. **E**) Untreated *wt* and *rd1* retina were compared to drug-treated retina as in A. Untr. *wt*: 8; Untr. *rd1*: 13; INO1001 *rd1*: 13; JA2131 *rd1*: 8; INO1001+JA2131 *rd1*: 10; 8-Br-ADPR *rd1*: 12. Statistical testing: One-way ANOVA with Tukey’s multiple comparison *post hoc* test comparing *rd1* explant cultures. Error bars represent SD; ns = *p* > 0.05; * = *p* < 0.05; ** = *p* < 0.01; *** = *p* < 0.001; **** = *p* < 0.0001. INL = inner nuclear layer, GCL = ganglion cell layer; scale bar = 50 µm.

To confirm the efficacy of PARP inhibitors, we performed an *in situ* PARP activity assay and an immunostaining for PAR, *i.e*. a staining for the product of PARP activity. While PARP activity and PAR-positive cells were infrequent in *wt* retina when compared with *rd1* retina, treatment with INO1001 significantly reduced both PARP activity and PAR accumulation in the *rd1* ONL (Figure 2B-E; Table S2A, B; dose-response curve shown in Figure S1A). However, the PARG inhibitor, JA2131, did not affect PARP activity and PAR in *rd1* ONL (Figure 2B-E; Table S2A, B). In contrast, PARP activity and PAR positive cells in *rd1* ONL were significantly reduced after treatment with the combination of INO1001 and JA2131, as well as after single treatment (*i.e.* monotherapy) with 8-Br-ADPR (Figure 2B-E; Table S2A, B). Also in *rd1*Cngb1^−/−^* retinas, which lack functional cGMP-activated CNG-channels, both Olaparib and INO1001 significantly reduced PARP activity and PAR generation (Figure S2A-D), suggesting that PARP activity was at least in part independent of CNG-channel function. Overall, this data showed that both INO1001 and 8-Br-ADPR efficiently reduced PARP activity, while inhibition of PARG with JA2131 did not.

### PARP, PARG, and TRPM2 regulate calpain activity and calpain-2 activation

In a very recent study, we found the activity of Ca^2+^-dependent calpain-type proteases to likely be controlled by PARP activity [23]. To further study how PARP, PARG, and TRPM2 inhibition impacted calpain activity, we first used a general *in situ* assay that can resolve enzymatic activity of calpains in individual retinal cells [37]. Calpain activity was rather low in *wt* retina when compared with *rd1* retina (Figure 3A; Table S2C). Treatments with INO1001, JA2131, INO1001 combined with JA2131, and 8-Br-ADPR all led to a significant reduction of the numbers of ONL cells showing calpain activity (Figure 3B; Table S2D; dose-response curve shown in Figure S1A, B, C, respectively). Remarkably, neither Olaparib nor INO1001 treatment could reduce calpain activity in *rd1*Cngb1^−/−^* photoreceptors, *i.e.* in photoreceptors in which CNG-channels are dysfunctional (Figure S3A, B). Since calpain activity in *rd1* single-mutant retina was significantly higher than in the *rd1*Cngb1^−/−^* situation, this outcome suggested that calpain activation was in part caused by CNG-channel independent Ca^2+^-influx. Moreover, in the *rd1*Cngb1^−/−^* double mutant retina calpain activity was apparently triggered independent of PARP.

**Figure 3.**
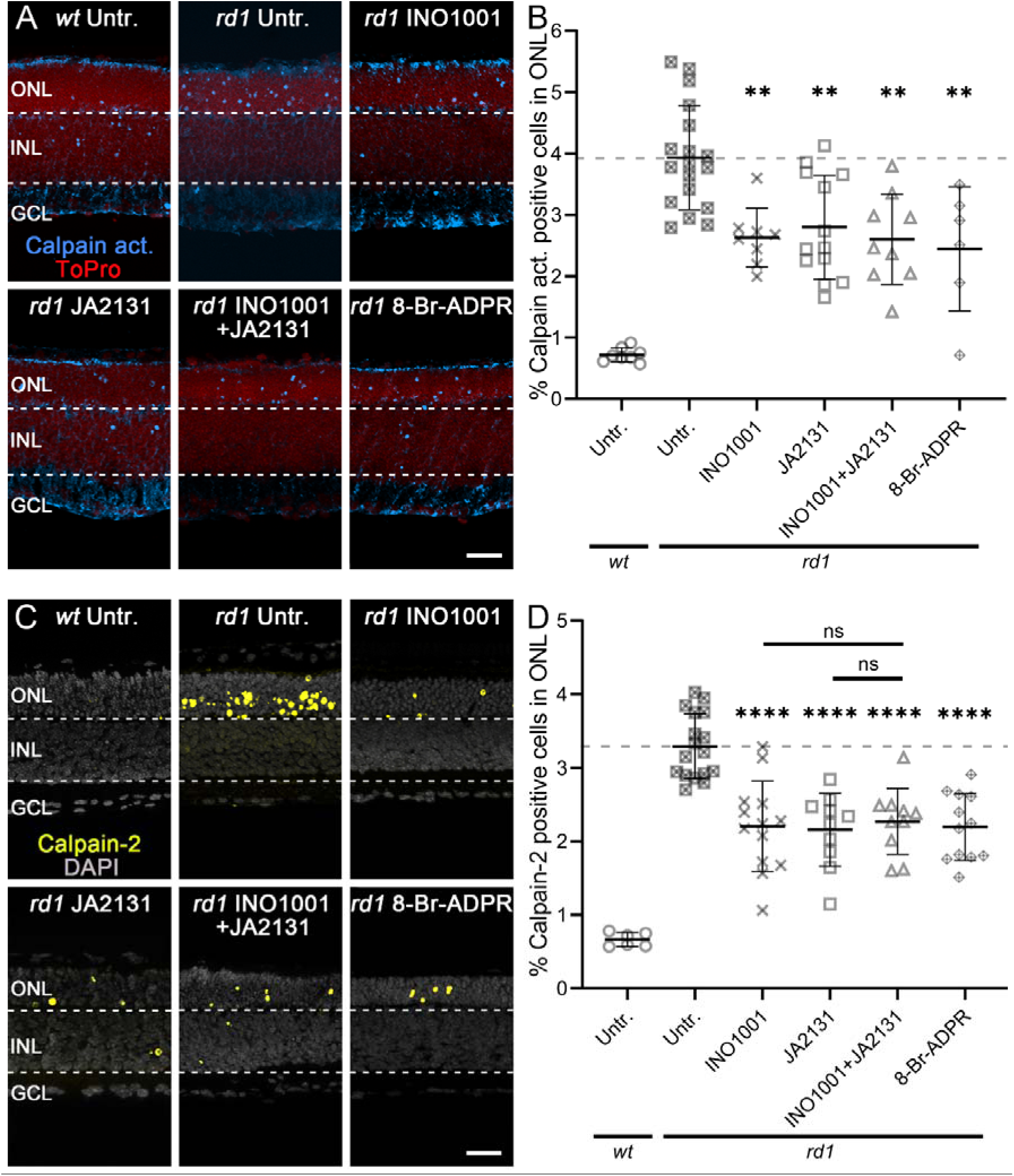
Inhibition of PARP-signalling decreases calpain activity and calpain-2 activation. **A**) Calpain activity assay (blue) was performed on wild-type (*wt*) and *rd1* retinal explant cultures. ToPro (red) was used as a nuclear counterstain. Untreated (Untr.) *wt* and *rd1* retina were compared to retina treated with INO1001, JA2131, INO1001 combined with JA2131, and 8-Br-ADPR. Note the high number of cells displaying calpain activity in the outer nuclear layer (ONL). **B**) Scatter plot showing percent calpain activity positive cells in the outer nuclear layer (ONL). Untr. *wt*: 8; Untr. *rd1*: 18; INO1001 *rd1*: 8; JA2131 *rd1*: 13; INO1001+JA2131 *rd1*: 8; 8-Br-ADPR *rd1*: 6. **C**) Immunostaining for activated calpain-2 (yellow) was performed using wild-type (*wt*) and *rd1* retinal explant cultures. DAPI (grey) was used as a nuclear counterstain. Untreated *wt* and *rd1* retina were compared to retina treated with INO1001, JA2131, 8-Br-ADPR, or a combination of INO1001 and JA2131. **D**) Scatter plot showing the percentage of ONL cells displaying calpain-2 activation. Untr. *wt*: n=6; Untr. *rd1*: 17; INO1001 *rd1*: 13; JA2131 *rd1*: 10; INO1001+JA2131 *rd1*: 10; 8-Br-ADPR *rd1*: 12. Statistical testing: One-way ANOVA and Tukey’s multiple comparison post hoc test; significance levels: ** = *p* < 0.01; **** = *p* < 0.0001; error bars represent SD; INL = inner nuclear layer, GCL = ganglion cell layer; scale bar = 50 µm.

Among the calpain-type proteases, calpain-2 appears to be the isoform that is most important for neurodegenerative processes [38] and immunolabelling for activated calpain-2 may provide for a snapshot in time that allows to infer changes in intracellular Ca^2+^ levels. Similar to total calpain activity, the number of cells in the ONL of *wt* retina displaying calpain-2 activation was rather low when compared with *rd1* retina (Figure 3C; Table S2D). In *rd1* retina, the drugs INO1001, JA2131, and ADPR, individually, significantly reduced calpain-2 activation (Figure 3D; Table S2D). A combined inhibition of PARP and PARG with INO1001 and JA2131 did not produce an additional synergistic effect (Figure 3D; Table S2D), indicating that both enzymes belonged to the same calpain-2 activating pathway. However, similar to what was seen for general calpain activity, in *rd1*Cngb1^−/−^* double-mutant retina the PARP inhibitors Olaparib and INO1001 could not reduce calpain-2 activation (Figure S3C, D).

### Inhibition of PARP, PARG, TRPM2 alters HDAC activity and reduces cell death

The activity of HDAC was previously suggested to be contributing to photoreceptor degeneration [39, 40]. We investigated a possible link to PARP-, PARG-, and TRPM2-related signalling using a general HDAC *in situ* activity assay that non-selectively detects activity of isoforms belonging to all four classes of HDACs. In *rd1* retina, the number of HDAC activity positive cells in the ONL was higher compared to *wt* (Figure 4A; Table S3A). Unexpectedly, INO1001 significantly increased overall HDAC activity in *rd1* ONL, while treatment with JA2131, INO1001+JA2131, and 8-Br-ADPR did not change the numbers of HDAC activity positive cells, as compared to untreated *rd1* cultures (Figure 4B; Table S3A). When compared to the monotherapies of INO1001 and JA2131, combined treatment with INO1001+JA2131 significantly reduced HDAC activity (Figure 4B; Table S3A).

**Figure 4.**
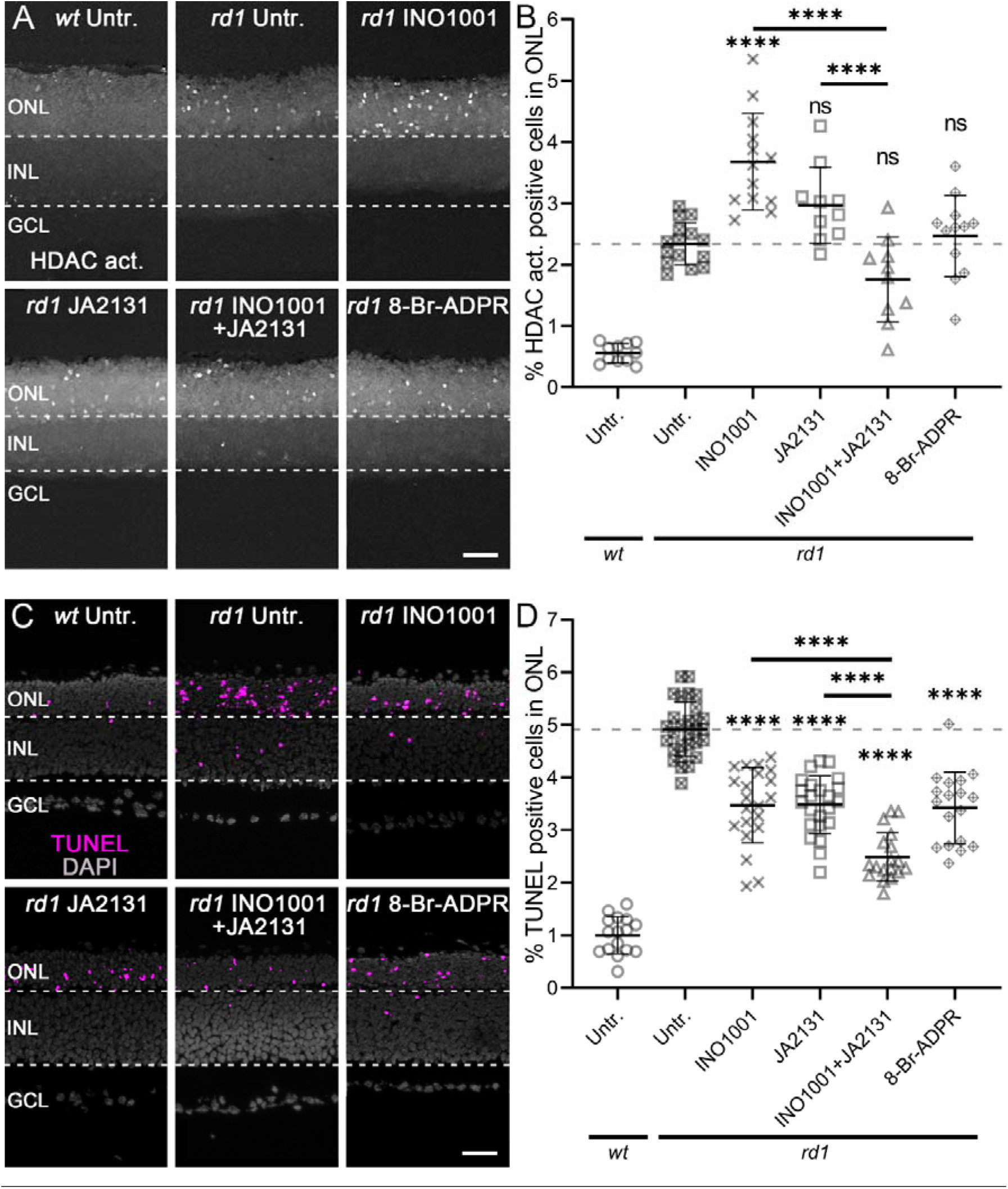
HDAC activity, and photoreceptor cell death after the inhibition of PARP-signalling. A) HDAC activity assay (white) was performed in *rd1* and *wt* retinal explant cultures. Untreated *rd1* and *wt* retina were compared to *rd1* retina treated with either INO1001, JA2131, INO1001+JA2131, or 8-Br-ADPR. **B**) Scatter plot showing percent displaying HDAC activity. Untr. *wt*: 11; Untr. *rd1*: 15; INO1001 *rd1*: 13; JA2131 *rd1*: 10; INO1001+JA2131 *rd1*: 10; 8-Br-ADPR *rd1*: 12. **C**) TUNEL assay labelling dying cells (magenta) in *rd1* and *wt* retinal explant cultures. DAPI (grey) was used as a nuclear counterstain. Untreated *wt* and *rd1* retinas were compared to retinas treated with compounds as in A. Untr. *wt*: 16; Untr. *rd1*: 29; INO1001 *rd1*: 21; JA2131 *rd1*: 23; INO1001+JA2131 *rd1*: 18; 8-Br-ADPR *rd1*: 18. **D**) Scatter plot showing the percentage of TUNEL positive cells. Note the significant elevation of HDAC activity caused by INO1001 alone and the synergistic protective effect on TUNEL positive cells when INO1001 and JA2131 treatments were combined. Statistical testing: One-way ANOVA and Tukey’s multiple comparison *post hoc* test; significance levels: ns = *p* > 0.05; **** = *p* < 0.0001; error bars represent SD; INL = inner nuclear layer, GCL = ganglion cell layer; scale bar = 50 µm.

To confirm the link between PARP-signalling and photoreceptor degeneration, the TUNEL assay was used to quantify the numbers of dying cells in the ONL. In *wt* retinal cultures, the numbers of TUNEL positive cells in the ONL was significantly lower than in its *rd1* counterpart (Figure 4C, Table S3B). Both INO1001 and JA2131 significantly decreased photoreceptor cell death in *rd1* cultures. Remarkably, INO1001 combined with JA2131 reduced the numbers of TUNEL positive cells in the ONL even further, when compared with untreated *rd1*, and monotherapies of INO1001 or JA2131 (Figure 4D, Table S3B). This was surprising since PARP and PARG would be expected to take part in the same pathway [41]. To confirm the synergistic effect observed, we combined JA2131 with Olaparib treatment in *rd1* retinal cultures. However, this combination failed to show the synergistic effect seen with INO1001 and JA2131 (Figure S4A, B). Moreover, in *rd1*Cngb1^−/−^* double-mutant mice, INO1001 treatment reduced photoreceptor cell death when Olaparib did not (Figure S5A, B), strongly suggesting a difference in the mode of action or the targets for these two compounds.

### Enzyme activity patterns are altered by interventions targeting PARP-signalling

To compare the various enzymatic and cell death markers with each other, we normalized the experimental datasets by linear scaling, such that the lowest values were set to zero while the highest values were set to one. In *wt* retinas, all enzyme activity and cell death markers were generally low when compared with the *rd1* untreated groups (Figure 5A). In untreated *rd1*, the number of ONL cells showing HDAC activity was lower than calpain-2 activation, calpain activity, PARP activity, and PAR positive cells. Photoreceptor cell death was significantly reduced by monotherapies of INO1001, JA2131, and 8-Br-ADPR. However, INO1001 led to relatively high HDAC activity, while JA2131 caused high PARP activity and PAR accumulation compared to other markers (Figure 5A). The combined treatment with INO1001 and JA2131 synergistically reduced photoreceptor degeneration and all related markers, relative to monotherapies with INO1001 or JA2131. Although also 8-Br-ADPR led to relatively low levels for all enzyme activity markers, it did not decrease cell death as much as INO1001/JA2131 combination therapy (Figure 5A). The relative increase in overall HDAC activity caused by INO1001, but not by any of the other drug treatments, was remarkable and could indicate an activation of Sirtuin-type HDACs.

**Figure 5.**
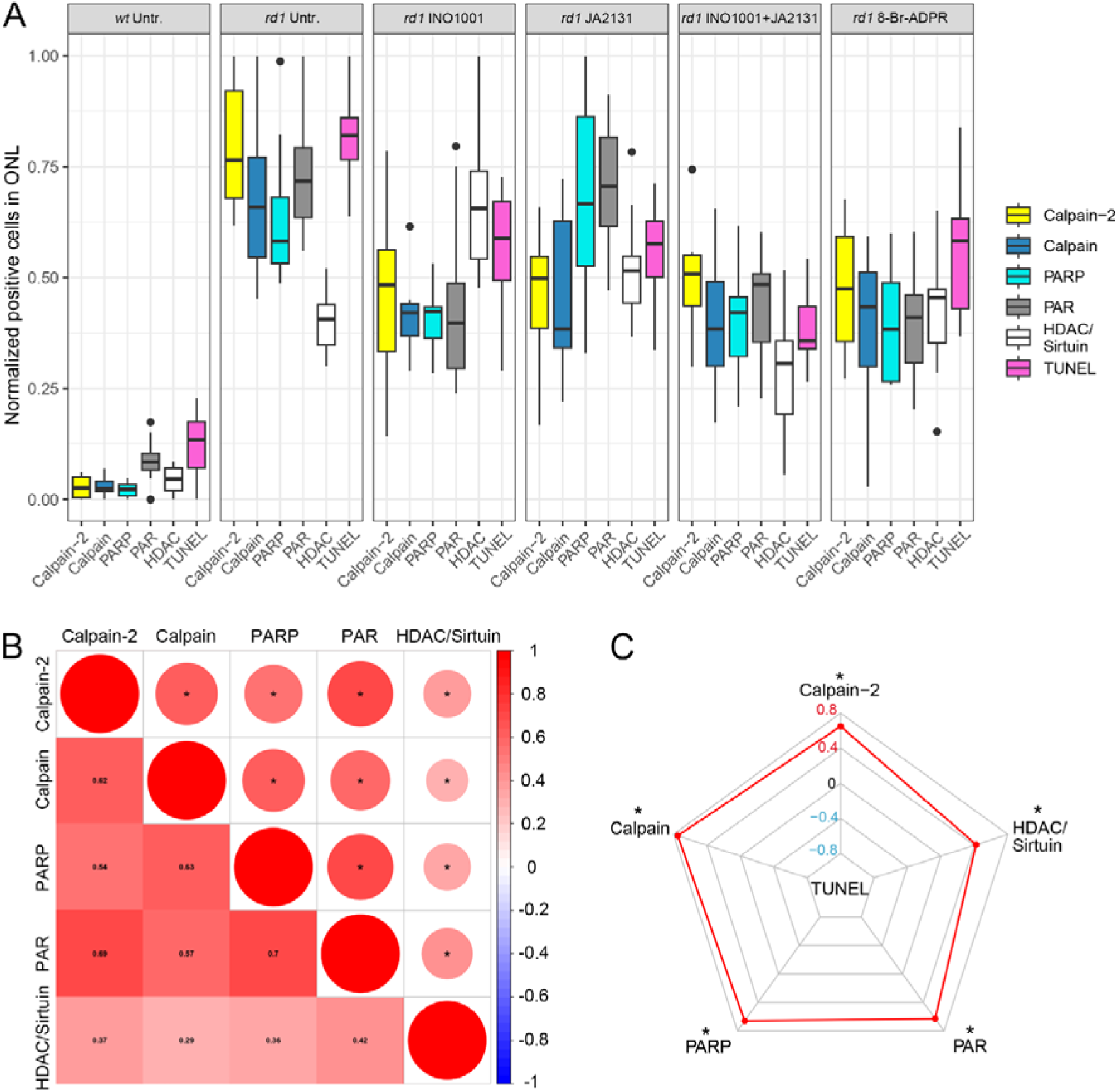
Enzymatic signatures for *rd1* photoreceptor degeneration and Spearman analysis. **A**) Comparison of enzymatic markers across different experimental treatments. Normalized cell numbers positive for TUNEL (magenta), calpain activity (calpain, blue), activated calpain-2 (yellow), PARP activity (PARP, cyan), PAR (black), and HDAC/Sirtuin activity (white). **B**) Spearman analysis of enzymatic markers in photoreceptors (calpain activity, calpain-2, PARP activity, PAR, HDAC). The asterisks in circles show statistical significance, numbers in squares present the *R^2^*. **C**) Radar plot for Spearman analysis revealing the correlation between different enzymatic markers and the TUNEL assay. Note the comparatively weak association between HDAC activity and cell death (TUNEL).

To better understand these activity patterns triggered by inhibition of PARP, PARG, or TRPM2, we performed a Spearman analysis (Figure 5B), including all datasets. General calpain activity, calpain-2 activation, PARP activity, PAR, and HDAC were positively correlated with each other (*p* < 0.05). A radar plot was used to show the relationship of cell death and enzymatic signatures after Spearman analysis (Figure 5C). In the *rd1* ONL, this illustrated a positive correlation between cell death (TUNEL positive cells) calpain activity, calpain-2 activation, PARP activity, and PAR, but less so with HDAC activity (*p* < 0.05, Figure 5C). Taken together, these analyses indicated that calpain activity, calpain-2, PARP, PAR, and HDAC activity were all connected to cell death. Then again, the association of HDAC activity with cell death was not as strong as for the other four markers, suggesting that among the various HDAC isoforms/classes there could also be such that might have protective effects.

## DISCUSSION

In the *rd1* mouse retina we previously found PARP to regulate calpain activity, promoting photoreceptor degeneration [9]. However, the mechanism of this PARP-dependent regulation of calpain was still unclear. Our present work proposes that PARP controls calpain activity likely through PARG-induced ADPR formation and TRPM2-associated Ca^2+^ influx. The blockage of either PARP, PARG, or TRPM2 was found to be neuroprotective. Notably, the combined therapy with the PARP inhibitor INO1001 and the PARG inhibitor JA2131 produced a synergistic therapeutic effect in *rd1* retina. These data suggest not only PARP, but also PARG and TRPM2 as therapeutic targets for the treatment of IRD. Overall, our study indicates PARP-dependent Ca^2+^ regulation to be detrimental to photoreceptor viability.

### PARP-dependent Ca^2+^ overload in photoreceptors

In the present study, we investigated the roles of PARP, PARG, and TRPM2 in photoreceptor cell death (Figure 6). PARP activity is connected to the *rd1* mutation in the *Pde6b* gene, likely via increased photoreceptor cGMP-levels (Figure 6) [5, 23]. Previously, we found the PARP inhibitor Olaparib to reduce the enzymatic activity of Ca^2+^-dependent calpain-type proteases and to slow down the progression of *rd1* photoreceptor degeneration [9]. Here, we find the alternative PARP inhibitor, INO1001, to show similar neuroprotective effects, consolidating the idea that PARP plays an important role in photoreceptor cell death. PARP catalyses the covalent attachment of PAR polymers onto itself by using NAD^+^ as a donor of ADPR units [12]. Covalently attached PAR can be hydrolysed to free PAR polymers or ADPR monomers by PARG, which possesses both endoglycosidic and exoglycosidic activities, and PAR hydrolase (ARH3), which also has PARG activity [12]. Overactivation of PARP may lead to NAD^+^ depletion [4], which will decrease a photoreceptors capacity to produce ATP. At the same time, increased generation of ADPR by PARP and PARG and consequent activation of TRPM2 will drive up Ca^2+^ influx [14, 42]. Consequently, ATP consumption for Ca^2+^ extrusion via the ATP-driven plasma membrane Ca^2+^-ATPase (PMCA) rises (Figure 6). Overall, a PARP-dependent decreased ATP production combined with increased consumption is likely to be strongly detrimental for photoreceptor cell viability [43].

**Figure 6.**
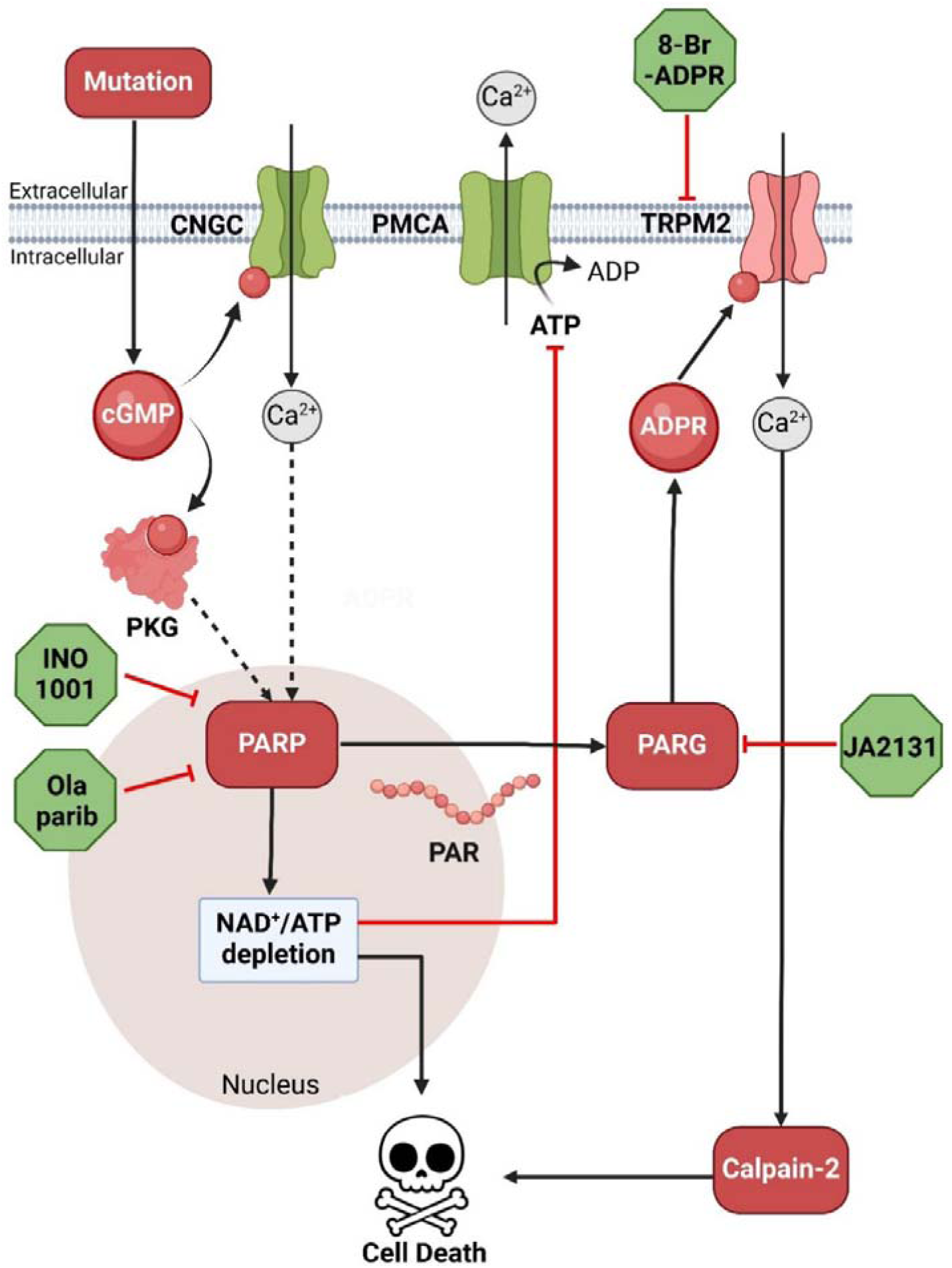
PARP-signalling and experimental interventions in cGMP-dependent *rd1* degeneration. In *rd1* photoreceptors, the *Pde6b* mutation induces cGMP accumulation, which activates both the cyclic-nucleotide-gated channel (CNGC) and protein kinase G (PKG). Via yet unknown mechanisms excessive PKG activity may result in DNA damage, which likely leads to poly (ADP-ribose)-polymerase (PARP) activation. On the one hand, poly (ADP-ribose) (PAR) generated by PARP may be cleaved by poly (ADP-ribose) glycohydrolase (PARG) into ADP-ribose (ADPR). ADPR can directly open transient receptor potential cation channel (TRPM2) leading to Ca^2+^ influx. On the other hand, PARP activation may entail a depletion of NAD^+^ and ATP. Since ATP is required for plasma membrane Ca^2+^ ATPase (PMCA)-mediated Ca^2+^ extrusion, loss of ATP and TRPM2 opening, in concert, lead to increased intracellular Ca^2+^ levels and excessive activation of calpain-type proteases. Eventually, the activities of calpain-2, PARP, and intracellular Ca^2+^ overload promote photoreceptor cell death.

When we used the PARG inhibitor JA2131 [30] on *rd1* retinal explant cultures, calpain activity and calpain-2 activation were significantly reduced, suggesting that ADPR changes induced by PARG inhibition indeed also affected intracellular Ca^2+^ levels in *rd1* retina. The photoreceptor protective effect of PARG inhibition is in line with an earlier study which showed that a genetic deletion of PARG was neuroprotective [35]. The fact that both PARP and PARG inhibition reduced calpain activity – and by extension Ca^2+^-influx – hinted at TRPM2 as a likely candidate for causing Ca^2+^ overload in *rd1* photoreceptors. This idea was supported by the use of the TRPM2 ion channel antagonist 8-Br-ADPR [44], which significantly reduced the enzymatic activity of calpain. TRPM2 inhibition also decreased PARP activity suggesting a possible Ca^2+^-mediated feedback loop that regulated PARP activity. This is consistent with a recent study which found that Ca^2+^-levels regulated PARP activity and promoted photoreceptor degeneration [23]. It seems likely that an elevated intracellular Ca^2+^ concentration leads to activation of calpain type proteases. Among these, calpain-2 – activated by millimolar Ca^2+^-levels – is likely to be particularly detrimental to cell survival, while calpain-1 – activated by much lower micromolar Ca^2+^-levels – may have a protective role [45]. Future studies using cell-type specific, conditional genetic manipulations of calpain-1 and -2 may yield further insights on the interplay of these calpain isoforms with intracellular Ca^2+^ and PARP activity. Taken together, our data indicates that in cGMP-induced photoreceptor degeneration PARP regulates Ca^2+^-dependent calpain activity through ADPR mediated activation of TRPM2.

### Efficacy of INO1001 in hereditary retinal degeneration

PARP activity was previously connected to photoreceptor degeneration [34] and different PARP inhibitors, including BMN- 673, 3-aminobenzamide, Olaparib, and PJ34 have shown neuroprotective effects in various IRD models [9, 46, 47]. Compared to these drugs, the PARP inhibitor INO1001 [48] used here showed an improved efficacy in the *rd1* mouse model. Surprisingly, the combination of INO1001 with the PARG inhibitor JA2131 showed a synergistic effect in reducing photoreceptor cell death without further reducing PARP and calpain activity, when compared to individual treatments with either INO1001 or JA2131. This suggests that the activities of PARP and calpain may be partially independent from the execution of photoreceptor cell death.

To further investigate this effect, we combined Olaparib with JA2131, which, however, did not synergistically decrease photoreceptor cell death in *rd1*. These results indicate that the mode of action or the targets of INO1001 could be different from Olaparib. In support of this view, a recent article found that INO1001 and Olaparib differ significantly in their inhibitory profiles. Both compounds potently inhibit the PARP-1 isoform, yet INO1001 inhibits PARP-2 only weakly while Olaparib also strongly blocks this isoform [29]. Whereas PARP-1 likely accounts for more than 90% of total cellular PARP activity [49], PARP-2 may under certain conditions have neuroprotective effects [50, 51]. Moreover, PARP-2 activity may be specifically enhanced by PAR chains [52], an effect that is counteracted by the activity of PARG. In a situation where PARP-1 is efficiently inhibited by INO1001 but PARP-2 remains active the additional inhibition of PARG by JA2131 may drive a selective increase in PARP-2 activity. It is therefore plausible to think that the synergistic beneficial effect seen in the combined treatment with INO1001 and JA2131 may depend on such an increase in PARP-2 activity. This could also in part explain why a synergistic effect of INO1001 and JA2131 was seen in the cell death assay but not in general PARP activity. However, at this point we cannot exclude the possibility that INO1001 hits targets entirely unrelated to PARP, and that these may instead be responsible for the observed beneficial effects. Even though such a coincidence may seem improbable, in this case the synergistic effect of INO1001 and JA2131 would be related to two independent cell death promoting pathways.

Besides, in the *rd1*Cngb1^−/−^* retina, which lacks functional CNG-channels [26], Olaparib failed to show a protective effect, while INO1001 significantly reduced photoreceptor cell death also in the double-mutant retina. This outcome likewise points at different modes of action for Olaparib and INO1001 and could furthermore indicate that CNG-channels may be more directly linked to PARP activity, perhaps because of compartmentalization and specific structural organization in the photoreceptor outer segments.

### Relationship between PARP and HDAC activity

The HDAC family consists of 18 different mammalian isoforms divided into four classes based on their sequence similarity to yeast counterparts [53, 54]. Depending on the mechanism of deacetylation of lysine residues on protein substrates, HDACs are divided into zinc-dependent (Class I, II, IV) and NAD^+^-dependent (Class III) isoforms. The latter group is commonly referred to as Sirtuins or Sirtuin-type HDACs [55]. Zinc-dependent HDACs were found to promote photoreceptor degeneration in the *rd1* mouse model for IRD [21, 40], even though within this group of HDACs there may also be individual isoforms with neuroprotective properties [56]. For NAD^+^-dependent Sirtuin-type HDACs an overall neuroprotective role appears very likely [57].

Since PARP is a major consumer of NAD^+^ and since Sirtuins are very sensitive to changes in NAD^+^-levels [58, 59], Sirtuin activity is often inversely correlated to PARP activity [60]. In our study, administration of INO1001 strongly and significantly increased HDAC activity in photoreceptors, without further increasing cell death. This hints at the possibility that the increased levels of HDAC activity seen with INO1001 treatment could refer to increased Sirtuin activity. A possible explanation for this effect might be that PARP inhibition and the resultant decrease in NAD^+^ consumption had allowed for Sirtuins to become (or remain) active. Also other works studies have found links between PARP and Sirtuins, notably activation of the Sirtuin-1 isoform [61, 62], supporting a close relationship between PARP and Sirtuin-type HDACs. Curiously, an increase of HDAC/Sirtuin activity was not seen after treatment of *rd1* explant cultures with the PARP inhibitor Olaparib [40], again pointing at differences in the mode of action between INO1001 and Olaparib. Future studies may address these differences in drug action and their possible root causes in, for instance, in different drug targets.

## Conclusion

In IRD, the activities of calpain and PARP are both closely associated with photoreceptor degeneration. Our results now demonstrate that in cGMP-induced photoreceptor degeneration, PARP regulates calpain activity, likely via ADPR and TRPM2 induced Ca^2+^ influx. We found that not only PARP activity, but likewise the activities of PARG and TRPM2 can cause photoreceptor degeneration. Expanding this research to also include genetic modulation of PARP-signalling could further refine these therapeutic targets. Our study also shows that while the PARP inhibitor Olaparib potently reduced general PARP activity, INO1001 presented an overall better therapeutic efficacy in both *rd1* and *rd1*Cngb1^−/−^* retina. Corresponding *in vivo* studies in IRD animal models, using appropriate delivery vehicles and administration routes, may validate the neuroprotective effects observed here and could facilitate future clinical translation. Taken together, these results emphasize the importance of PARP and its associated pathways as attractive targets for future therapeutic interventions in IRD.

## ACKNOWLEDGEMENTS

The authors would like to thank Norman Rieger (from Institute for Ophthalmic Research, Eberhard-Karls-Universität Tübingen) for excellent technical assistance.

## AUTHOR CONTRIBUTIONS

Conceptualization, J.Y. and F.P.-D.; methodology, J.Y., L.K; software, J.Y., L.K.; validation, J.Y.; formal analysis, J.Y. QL.Y., and QX.Y.; investigation, J.Y., L.W.; data curation, J.Y.; writing-original draft preparation, J.Y.,L.K.; writing—review and editing, J.Y., KW.J., ZL.H., and F.P-D.; visualization, J.Y., QL.Y., and QX.Y.; supervision, F.P-D.; project administration, F.P-D.; funding acquisition, F.P-D. All authors have read and agreed to the published version of the manuscript.

---

**Figure S1.**
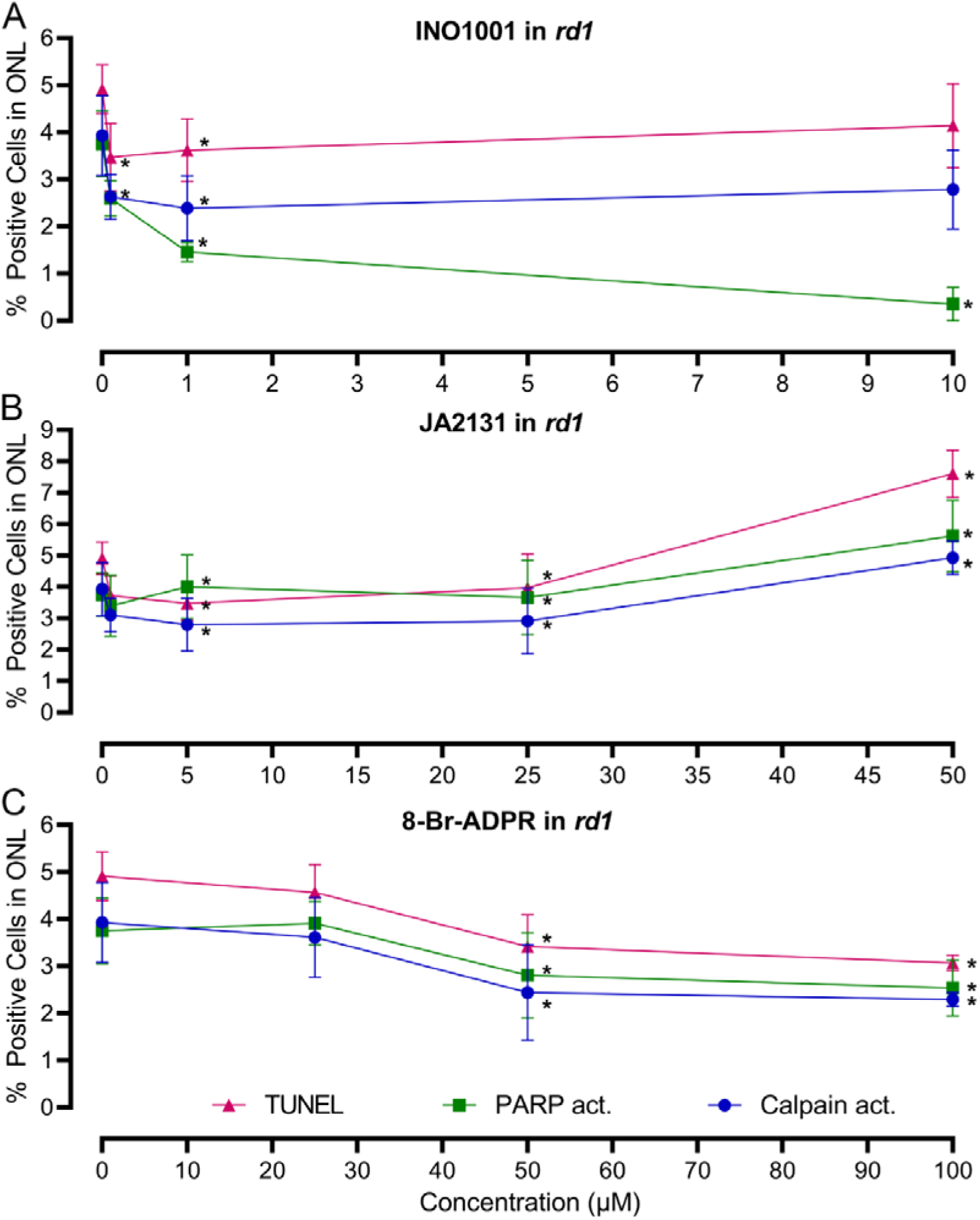
Dose-response curves for INO1001, JA2131, and 8-Br-ADPR. **A**) Different concentrations of INO1001 were tested in *rd1* mouse retinal explant cultures. In the outer nuclear layer (ONL), at concentrations of 0.1 µM and 1 µM, INO1001 significantly decreased the numbers of ONL cells displaying calpain activity, PARP activity, and cell death (TUNEL assay). **B**) Different concentrations of JA2131 tested in *rd1* explant cultures. In the ONL, at concentrations of 0.5 µM, 5 µM, and 25 µM, JA2131 significantly reduced ONL calpain activity, and cell death, as assessed by the TUNEL assay. **C**) Dose-response for 8-Br-ADPR in *rd1* explant cultures. 50 µM and 100 µM 8-Br-ADPR significantly reduced calpain activity, PARP activity, and cell death (TUNEL) in the ONL. Statistical significance was assessed using one-way ANOVA and Tukey’s multiple comparison post hoc test; significance levels: * = *p* < 0.05 error bars represent SD.

**Figure S2.**
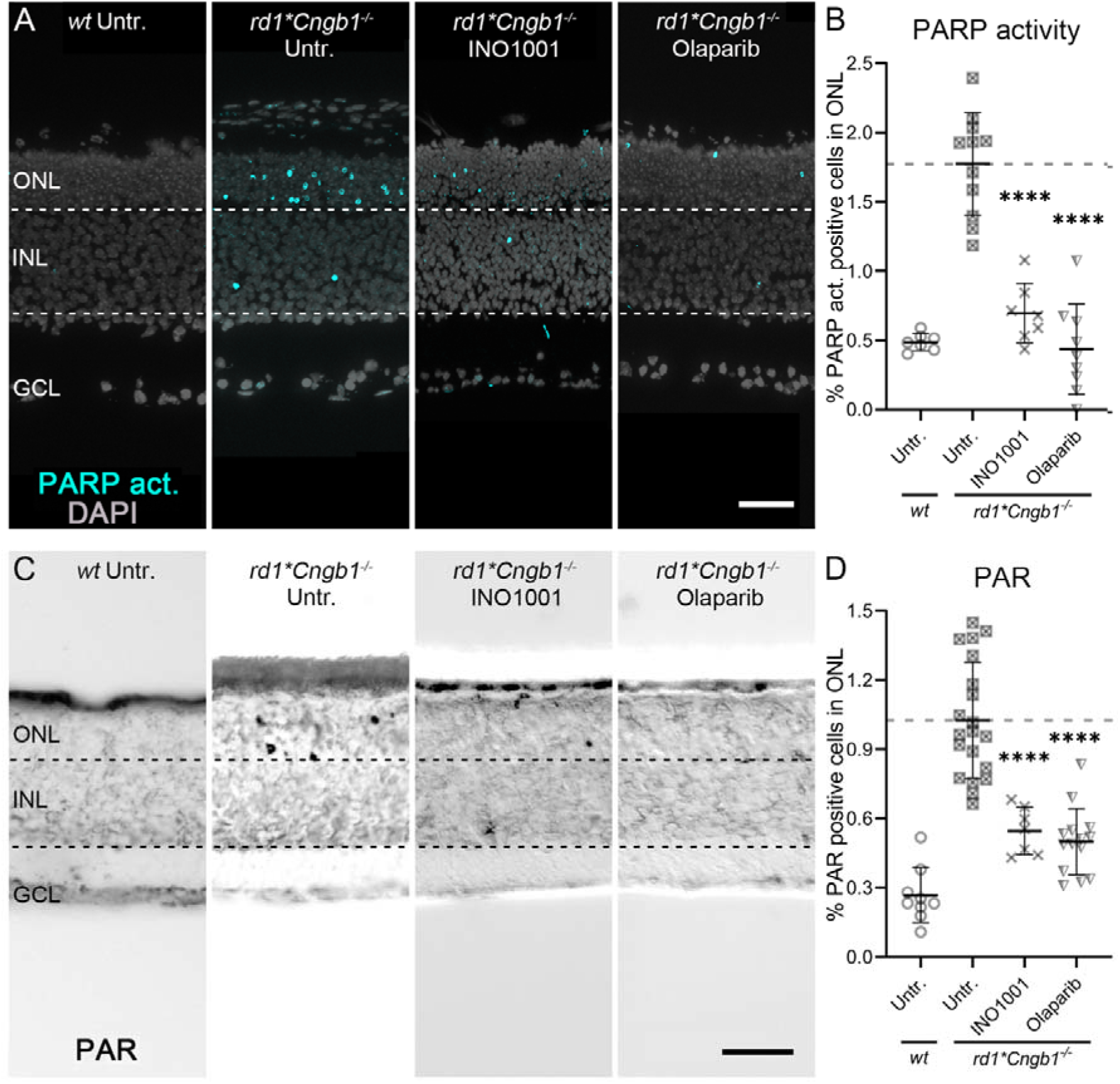
Effect of PARP inhibitors on *rd1*Cngb1^−/−^* retinal explant cultures. **A**) PARP activity assay (cyan) was performed in *rd1* and wild-type (*wt*) retinal explant cultures. DAPI (grey) was used as a nuclear counterstain. Untreated (Untr.) *rd1*Cngb1^-/-^*and *wt* retina were compared to *rd1*Cngb1^-/-^* retina treated with either INO1001 or Olaparib. **B**) Scatter plot showing the percentage of PARP activity positive cells in the outer nuclear layer (ONL). Untr. *wt*: 7; Untr. *rd1*Cngb1^-/-^*: 11; INO1001 *rd1*Cngb1^-/-^*: 7; Olaparib *rd1*Cngb1^-/-^*: 9. **C)** PAR staining (black) was performed in *rd1*Cngb1^-/-^*and *wt* retinal explant cultures. **D**) Untreated *wt* and *rd1*Cngb1^-/-^* retina were compared to drug-treated retina as in A. Untr. *wt*: 9; Untr. *rd1*Cngb1^-/-^*: 20; INO1001 *rd1*Cngb1^-/-^*: 7; Olaparib *rd1*Cngb1^-/-^*:14. Statistical testing: one-way ANOVA with Tukey’s multiple comparison post hoc test performed between *rd1* explant cultures. Error bars represent SD; **** = *p* < 0.0001. INL = inner nuclear layer, GCL = ganglion cell layer. Scale bar = 50 µm.

**Figure S3.**
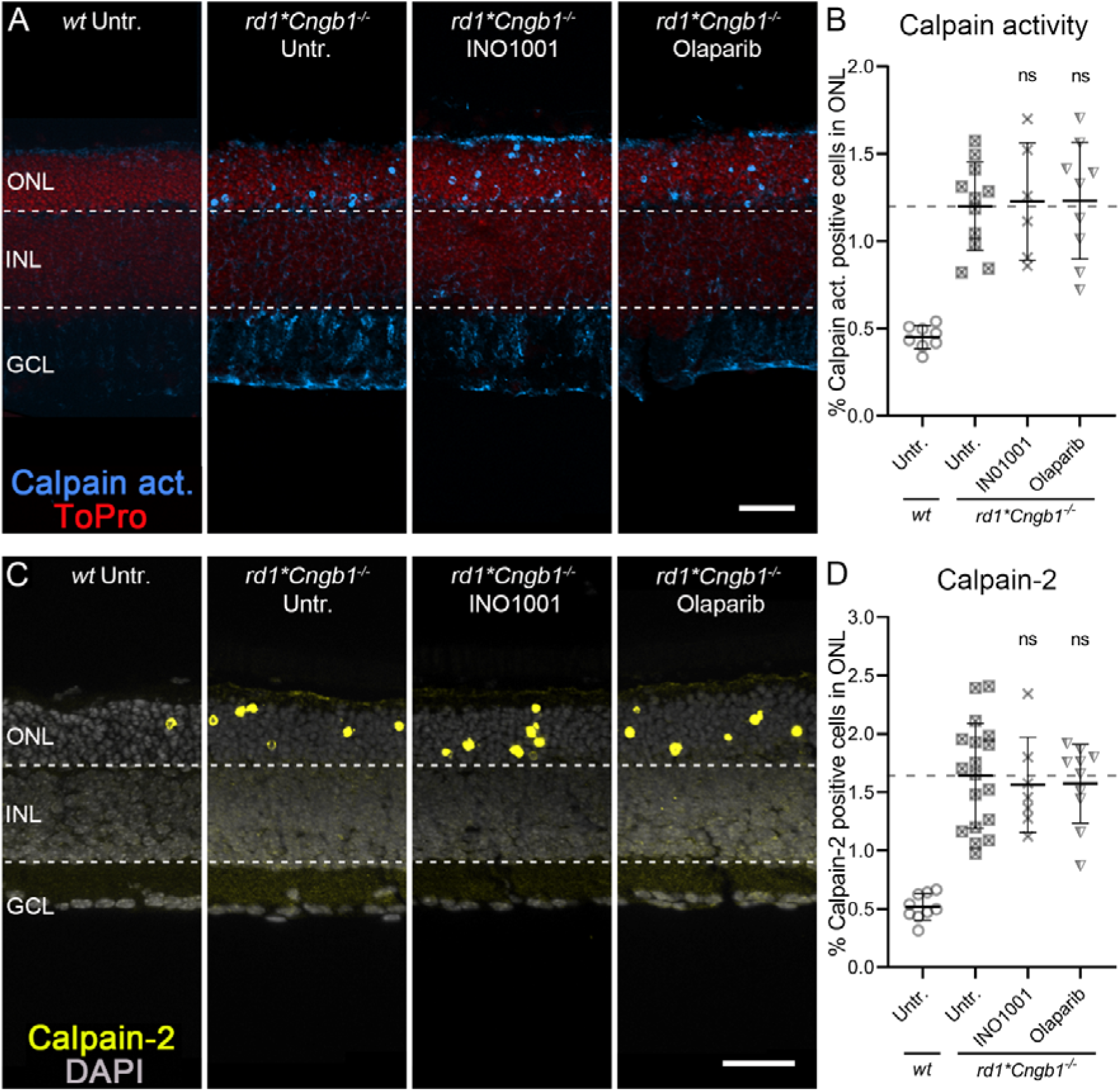
In *rd1*Cngb1^−/−^* retinal explant cultures PARP inhibitors do not affect calpain activity or calpain-2 activation. **A**) Calpain activity assay (blue) was performed on wild-type (*wt*) and *rd1*Cngb1^-/-^* retinal explant cultures. ToPro (red) was used as a nuclear counterstain. Untreated (Untr.) *wt* and *rd1*Cngb1^-/-^* retina were compared to retina treated with INO1001 and Olaparib. **B**) Scatter plot showing percent calpain activity positive cells in the ONL. Untr. *wt*: 8; Untr. *rd1*Cngb1^-/-^*: 11; INO1001 *rd1*Cngb1^-/-^*: 6; Olaparib *rd1*Cngb1^-/-^*: 9. **C**) Immunostaining for activated calpain-2 (yellow) was performed using wild-type (*wt*) and *rd1*Cngb1^-/-^* retinal explant cultures. DAPI (grey) was used as a nuclear counterstain. Untreated *wt* and *rd1*Cngb1^-/-^* retina were compared to retina treated with INO1001 and Olaparib. **D**) Scatter plot showing the percentage of ONL cells displaying calpain-2 activation. Untr. *wt*: 9; Untr. *rd1*Cngb1^-/-^*: 18; INO1001 *rd1*Cngb1^-/-^*: 7; Olaparib *rd1*Cngb1^-/-^*: 10. Statistical testing: one-way ANOVA with Tukey’s multiple comparison post hoc test performed between *rd1* explant cultures. Error bars represent SD; ns = *p* > 0.05. INL = inner nuclear layer, GCL = ganglion cell layer. Scale bar = 50 µm.

**Figure S4.**
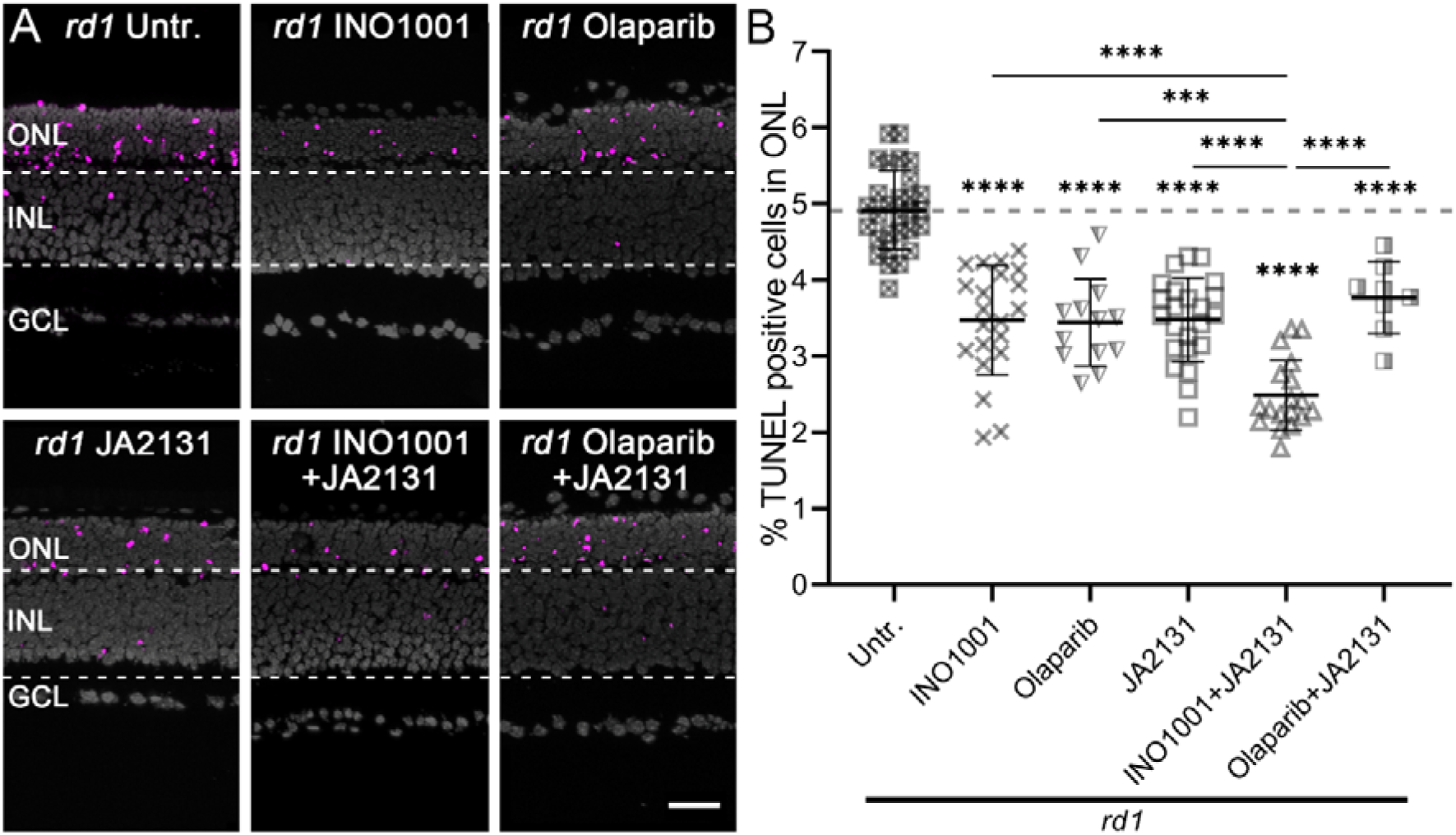
Inhibition of PARP/PARG reduces *rd1* photoreceptor cell death. **A**) TUNEL assay labelling dying cells (magenta) in *rd1* retinal explant cultures. DAPI (grey) was used as a nuclear counterstain. *rd1* untreated retina treated *rd1* retina were compared to *rd1* retina treated with INO1001, Olaparib, JA2131, INO1001+JA2131, and Olaparib+JA2131. Untr. *rd1*: 29; INO1001 *rd1*: 21; Olaparib *rd1*: 13; JA2131 *rd1*: 23; INO1001+JA2131 *rd1*: 18; Olaparib+JA2131 *rd1*: 8. **B)** Scatter plot showing percent displaying TUNEL positive cells. Note the synergistic effect of TUNEL positive cells triggered by INO1001+JA2131, but not by Olaparib+JA2131. Statistical testing: one-way ANOVA with Tukey’s multiple comparison post hoc test performed between *rd1* explant cultures. Error bars represent SD; *** = *p* < 0.001; **** = *p* < 0.0001. INL = inner nuclear layer, GCL = ganglion cell layer. Scale bar = 50 µm.

**Figure S5.**
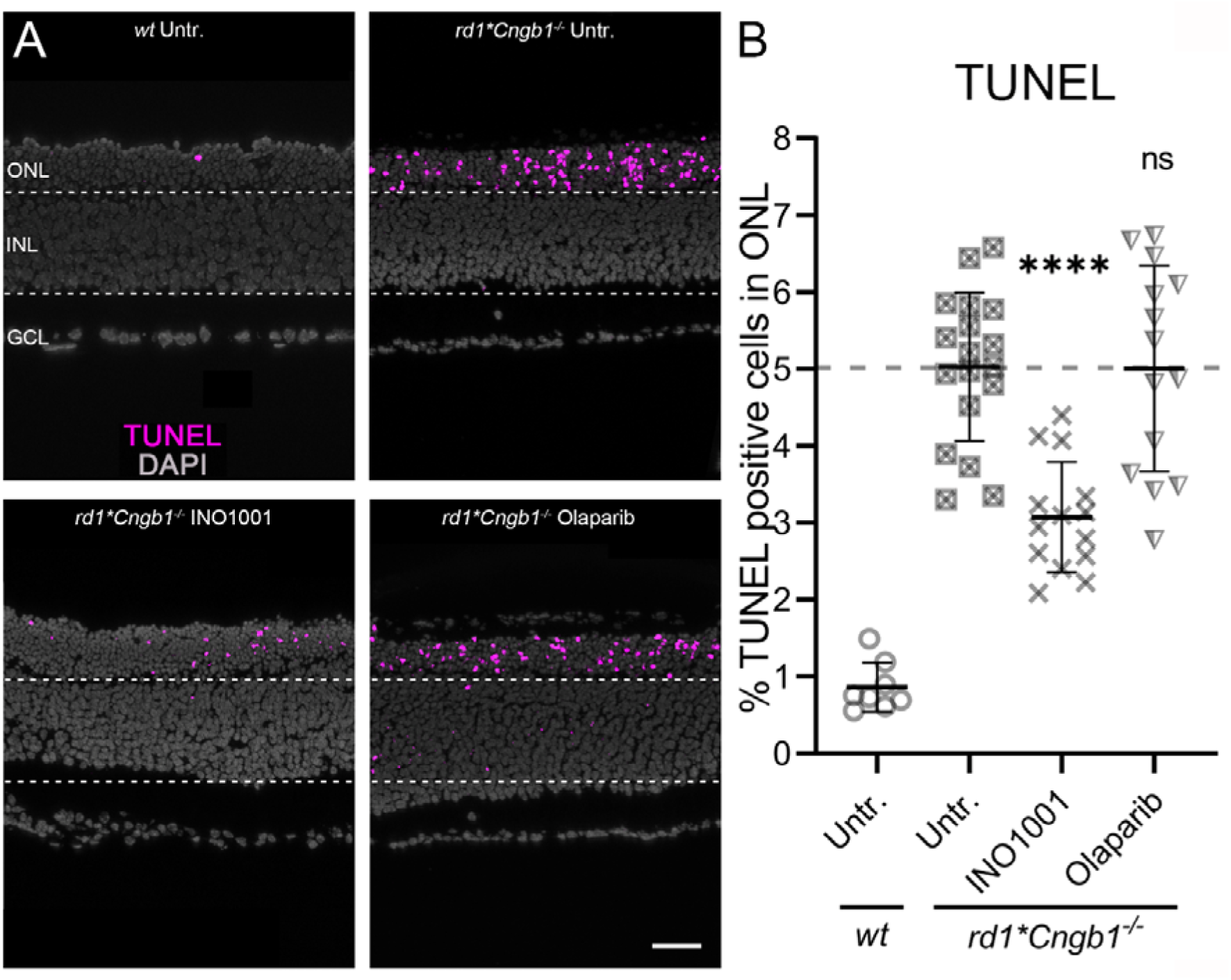
The PARP inhibitor INO1001 reduces *rd1*Cngb1^−/−^*cell death, when Olaparib does not. **A**) TUNEL positive cells (magenta) in *wt* and *rd1*Cngb1^-/-^* retinal cultures. rd1 retina was left either untreated (Untr.) or treated with INO1001 or Olaparib. **B**) Scatter plot showing percent displaying TUNEL positive cells. Statistical testing: One-way ANOVA with Tukey’s multiple comparison *post hoc* test performed between *rd1* explant cultures. Error bars represent SD; ns = *p* > 0.05; **** = *p* < 0.0001. INL = inner nuclear layer, GCL = ganglion cell layer. Scale bar = 50 µm.

**Supplemental Table 1.**
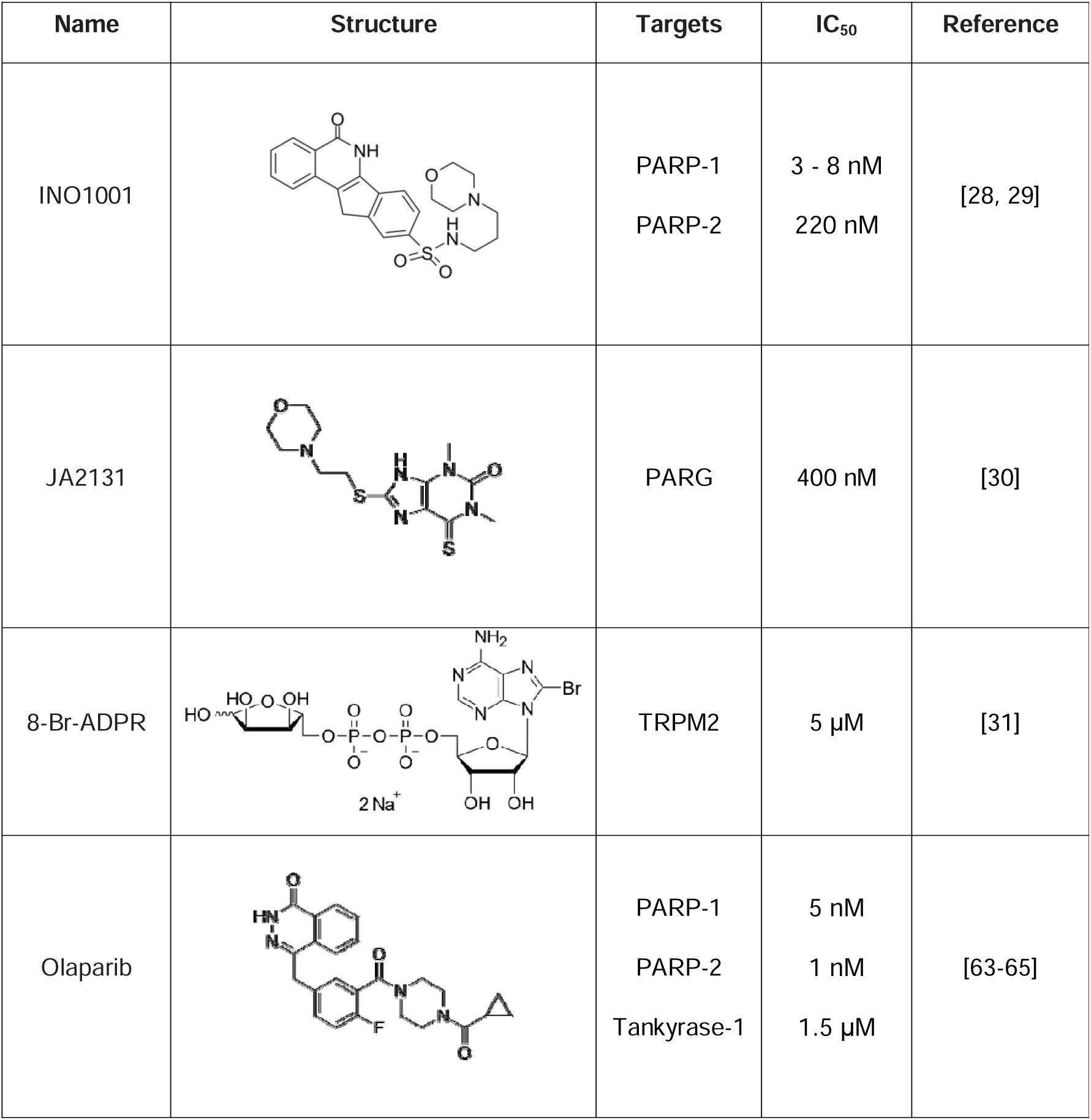
Information of compounds used in *rd1* explant cultures.

| Name | Structure | Targets | IC<sub>50</sub> | Reference |
| --- | --- | --- | --- | --- |
| INO1001 |  | PARP-1<br>PARP-2 | 3 - 8 nM<br>220 nM | [28, 29] |
| JA2131 |  | PARG | 400 nM | [30] |
| 8-Br-ADPR |  | TRPM2 | 5 μM | [31] |
| Olaparib |  | PARP-1<br>PARP-2<br>Tankyrase-1 | 5 nM<br>1 nM<br>1.5 μM | [63-65] |

**Supplemental Table 2.**
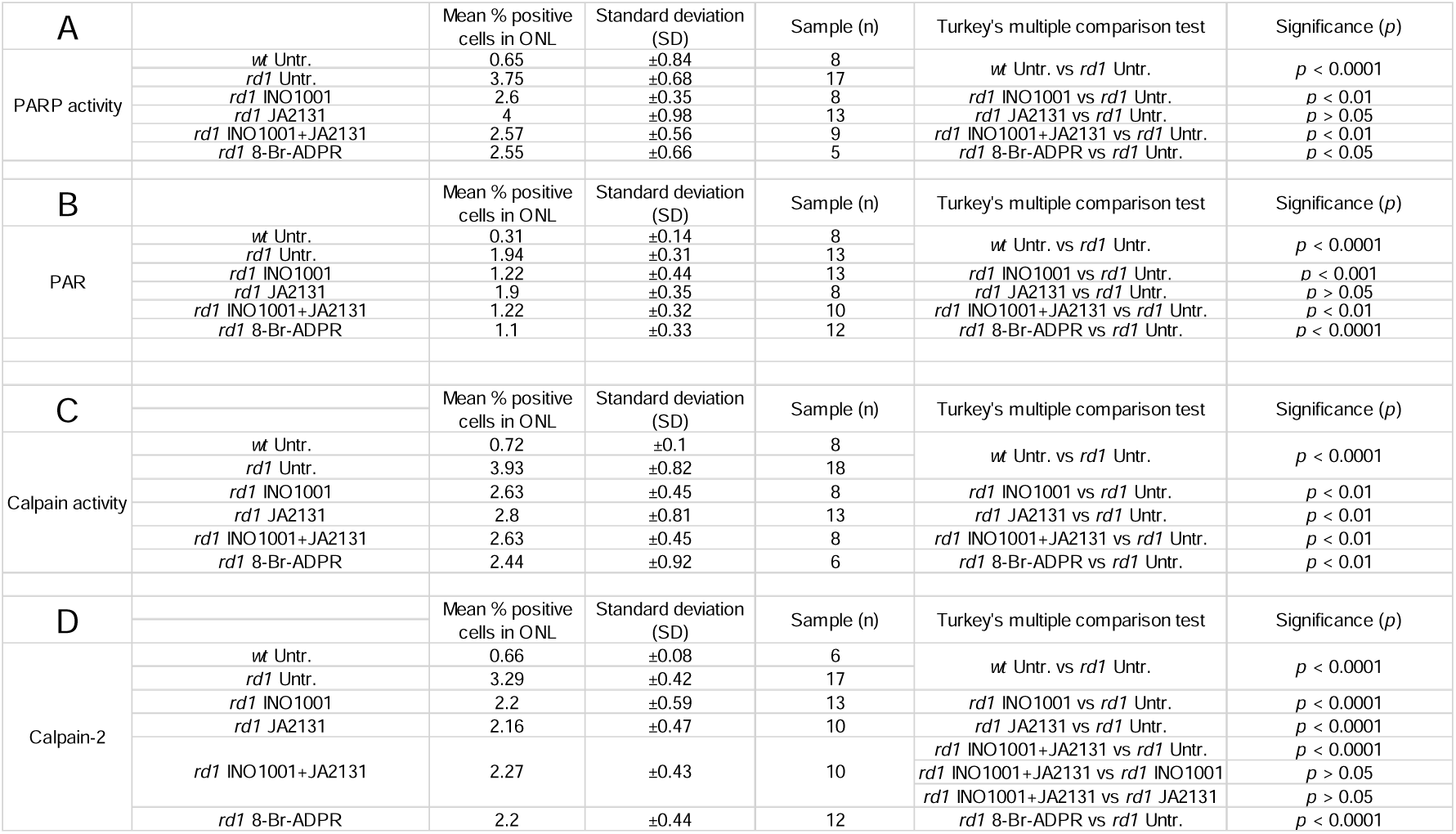
Quantification of calpain-2 activation, calpain activity, PARP activity, and PAR positive cells in the outer nuclear layer (ONL).

**Supplemental Table 3.**
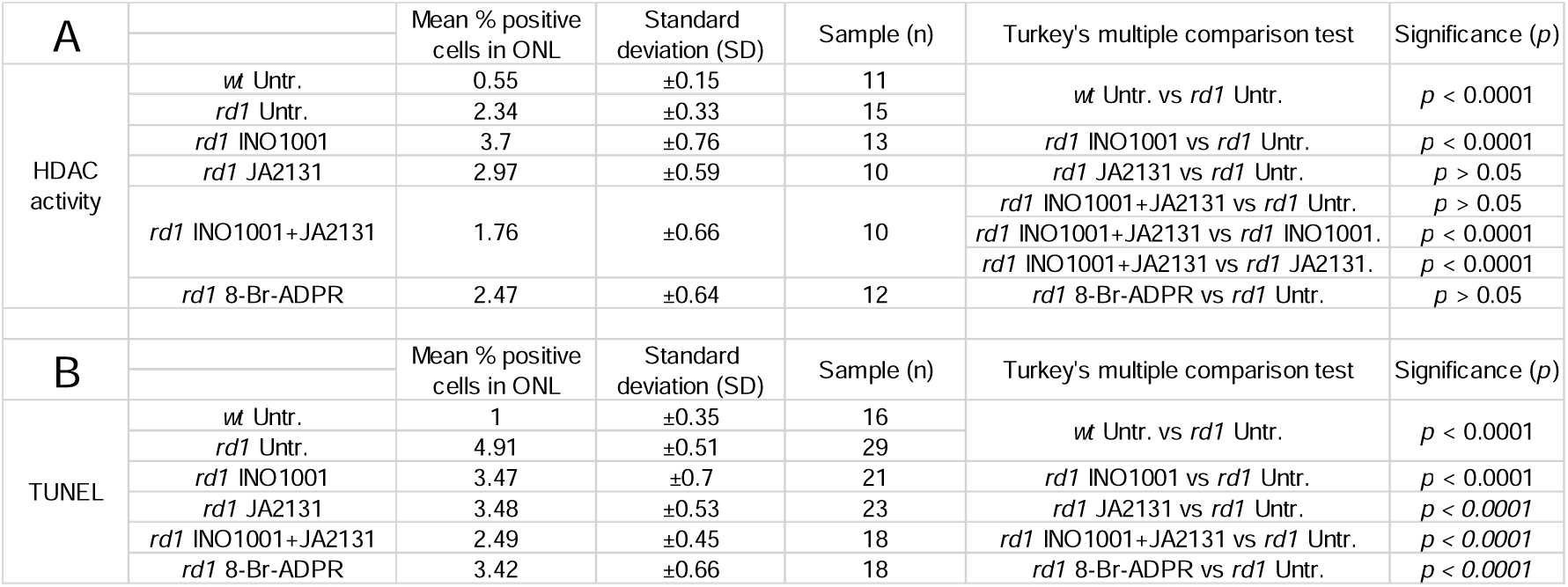
Quantification of HDAC activity and TUNEL positive, dying cells in the outer nuclear layer (ONL).

